# Behavioural and neural signatures across diverse cognitive demands in a multimodal electroencephalography-functional magnetic resonance imaging design

**DOI:** 10.64898/2026.05.24.727533

**Authors:** K. Hiromitsu, S. Chiyohara, T. Asai, A. Katayama, M. Wakabayashi, H. Imamizu

## Abstract

Efficient multimodal designs that capture differences across cognitive domains and variations in cognitive demand remain limited. In this study, we tested a compact framework with 58 healthy participants who completed multimodal electroencephalography (EEG) and functional magnetic resonance imaging (fMRI) sessions. The framework comprised two complementary batteries: the HCP-aligned multitask paradigm (HCP-mini), which integrates eight HCP-aligned cognitive tasks and rest within a single run, and an extended N-back task ranging from 0-back to 7-back. Designed to support broad cross-domain coverage and matched multimodal assessment, the two batteries captured the expected group-level behavioural structure across modalities. Behavioural performance exceeded chance levels or aligned with findings from previous studies in both EEG and fMRI. Descriptive intraclass correlation coefficient (ICC) analyses showed numerically higher within-modality run-to-run values than between-modality values. At the neural level, HCP-mini fMRI activation patterns closely recapitulated the canonical large-scale task organisation of the original HCP dataset, with corresponding task pairs showing the strongest spatial similarity. Together, these findings demonstrate a compact and efficient framework for multimodal characterisation of cognition across domains and graded cognitive demands.

## Introduction

Characterising the neural basis of cognition across multiple cognitive domains has become essential in cognitive neuroscience, particularly for distinguishing between task-general and task-specific neural processes (Cai et al., 2024; Cole et al., 2014; Cole et al., 2016). Carefully controlled tasks targeting specific functions have been highly informative for mapping regional task activations; however, they do not effectively reveal how task representations are organised across the brain. In contrast, studying multiple tasks within the same individual can help clarify the broader principles by which a single brain architecture supports diverse cognitive functions (Ito & Murray, 2022). Using the multidomain task battery developed by King et al. (2019), Ito and Murray (2022) showed that multitasking representations across the cortex are organised along a sensory-to-motor hierarchy. These considerations have motivated designs in which multiple cognitive tasks are measured under common acquisition conditions, directly comparing neural responses across tasks within individuals. Such designs are valuable for examining the relationships between cognitive functions, characterising differences in cognitive demand across tasks, and strengthening the estimation of participant-specific functional organisations (Gratton et al., 2020; Lynn et al., 2021; Poldrack et al., 2013). Although large-scale network architecture is broadly preserved across tasks, task-dependent modulation of functional connectivity shapes task-evoked activation patterns (Cole et al., 2021).

Multimodal neuroimaging provides further opportunities to characterise cognitive processes because different modalities capture complementary aspects of brain activity (Fleury et al., 2023). Electroencephalography (EEG) provides millisecond-scale temporal resolution, while functional magnetic resonance imaging (fMRI) provides spatially precise measurements of haemodynamic responses. Therefore, combining these modalities can offer a more comprehensive account of neural dynamics compared to using either modality alone (Debener et al., 2006; Warbrick, 2022).

Several studies and open datasets combining EEG and fMRI data have recently been reported. However, most studies have focused on a single experimental paradigm rather than on multiple cognitive tasks. Examples include perceptual decision confidence (Gherman & Philiastides, 2018), auditory and visual oddball paradigms (Walz et al., 2015), motor imagery neurofeedback (Lioi et al., 2020; Perronnet et al., 2017), and naturalistic viewing (Telesford et al., 2023). Other datasets acquired EEG and fMRI data in separate sessions using the same paradigm, such as the multimodal face recognition dataset (Wakeman & Henson, 2015) and recent naturalistic listening datasets (Momenian et al., 2024; Wang et al., 2025). Few studies or datasets included multiple cognitive tasks with multimodal measurements (Cha et al., 2026; Nakuci et al., 2023; Tsutsumi et al., 2026); however, these studies mostly relied on simultaneous EEG-fMRI acquisitions rather than separate-session measurements. Thus, multimodal designs that enable direct within-participant comparisons across multiple cognitive tasks remain limited, particularly when two modalities are acquired in separate sessions under matched protocols. This gap is notable because separate-session acquisition offers several methodological advantages, including improved EEG signal quality outside the MRI environment, direct comparison of neural responses across tasks within the same individual, and compatibility with representational analyses that relate time-resolved EEG signatures to spatially resolved fMRI patterns (Cichy & Oliva, 2020; Warbrick, 2022).

The Human Connectome Project (HCP) is one of the most influential frameworks for investigating task-evoked brain activity across multiple cognitive domains (Barch et al., 2013; Van Essen et al., 2013). Its task battery systematically organises functional imaging across domains, such as working memory, language, motor function, emotion, social cognition, incentive processing, and relational reasoning, thereby establishing a widely used reference framework for characterising domain-related activation patterns. Simultaneously, the original HCP task-fMRI design was developed to maximise efficiency while retaining the ability to identify robust task activations across a broad range of neural systems. Thus, the HCP task battery provides a comprehensive framework for examining task-evoked activity across multiple cognitive domains within a common experimental structure.

In the present study, we aimed to build on the HCP while introducing a more compact acquisition framework. First, we developed an HCP-aligned multitask paradigm (HCP-mini), in which eight cognitive tasks adapted from the HCP tasks and a resting-state condition were integrated within a single run of approximately 15 minutes. This compact design preserved the essential task structures of the original HCP paradigms while enabling direct comparison across cognitive domains within the same run (Barch et al., 2013; Cole et al., 2014). Second, we incorporated an extended N-back paradigm that parametrically manipulated the working memory load from 0-back to 7-back, with a resting-state condition, to capture the complementary axis of cognitive demand. The N-back task is a well-established paradigm for incrementally manipulating the working memory load, with systematic changes observed in both behavioural performance and neural activity as the load increases (Lamichhane et al., 2020). Thus, the HCP-mini and N-back paradigms were designed to capture two complementary dimensions of cognitive demands: cross-domain and within-task load variations (Figure 1). Therefore, in this study, we investigated whether a single-run HCP-aligned multitask framework, alongside a parametrically graded N-back task, could reproduce canonical behavioural signatures and task-related fMRI activation patterns comparable to those established in the original HCP framework across both cognitive domains and graded cognitive demand.

**Figure 1.**
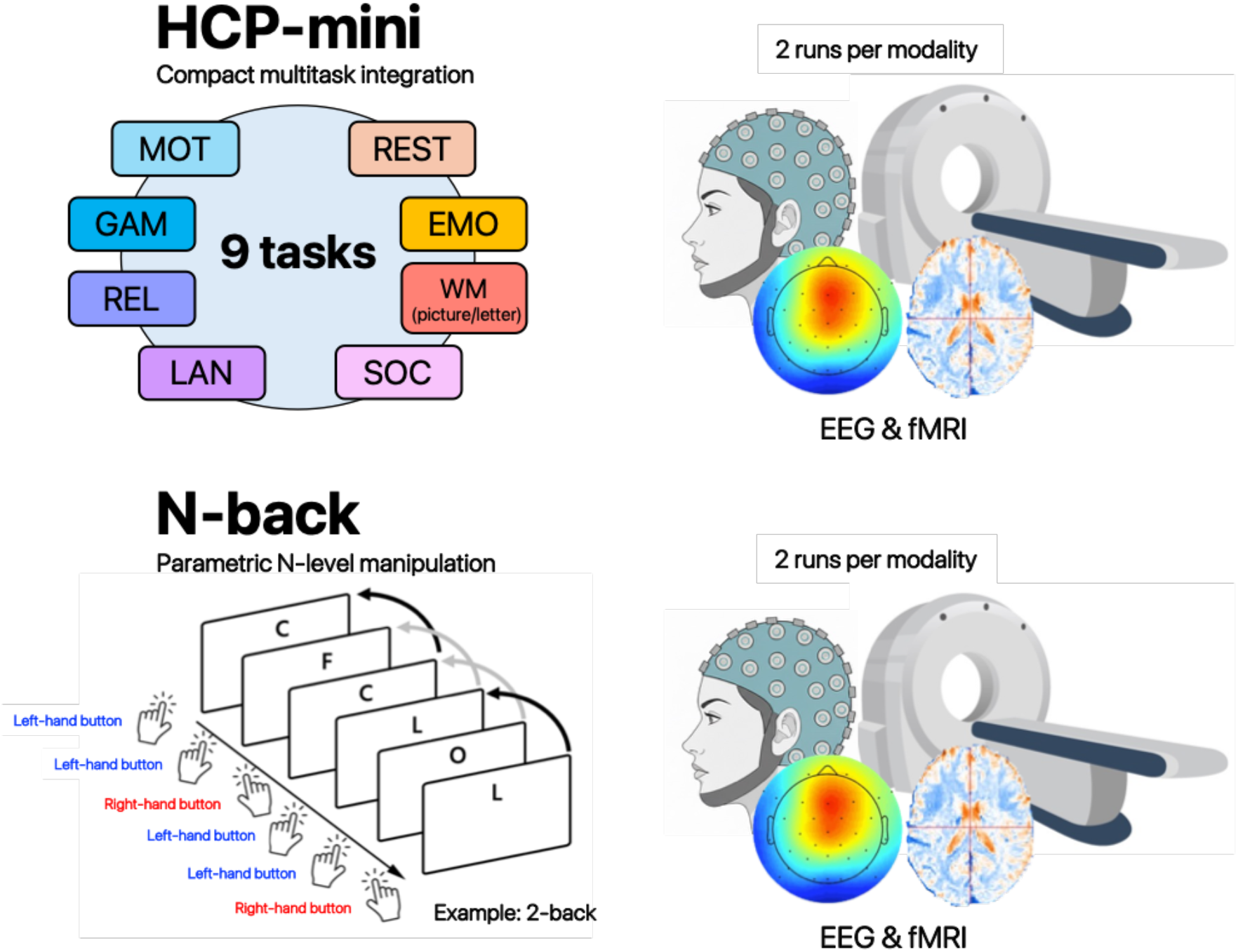
Overview of the experimental design and task structure in the study. The study includes two-task batteries acquired using EEG and fMRI. The first battery consists of a compact Human Connectome Project (HCP)-aligned multitask paradigm (HCP-mini), integrating eight cognitive tasks—emotion processing (EMO), social cognition (SOC), gambling (GAM), language (LAN), motor (MOT), relational processing (REL), and two working-memory variants (WM-picture and WM-letter)—together with a resting-state condition (REST) within a single run. The second battery is a parametric N-back task designed to manipulate working memory load across multiple levels. For each modality, two runs were conducted for each task battery. EEG and fMRI sessions are conducted on separate days using identical task structures, enabling multimodal comparisons of neural activity across cognitive paradigms.

## Methods

### Overview

This study evaluated whether a compact multitask acquisition strategy could reproduce established behavioural and fMRI signatures of cognitive demand using the HCP-mini and N-back paradigms. The HCP-mini paradigm prioritised broad cognitive coverage within a single run rather than retaining all explicit control conditions from the original HCP tasks. Accordingly, we adopted a cross-task comparative framework, in which activity for a given task was evaluated relative to the mean activity observed across all task conditions performed within the same run. This design was intended to characterise domain-dominant and domain-general activation profiles within a multitask framework rather than reproduce classical condition-specific contrasts (Cai et al., 2024; Cole et al., 2014; Fedorenko et al., 2013; Williams et al., 2022). Meanwhile, the extended N-back paradigm parametrically manipulated the working memory load to capture the complementary axis of cognitive demand. Its role was not merely to add another cognitive paradigm but to provide an independent dimension of cognitive demand. Specifically, whereas the HCP-mini task battery captured differences across cognitive domains, the N-back paradigm captured graded differences in cognitive demand within a single task. Notably, both paradigms shared a parallel nine-condition structure (eight active conditions plus rest), allowing cross-domain and within-task load variations to be examined within a parallel comparable framework. All tasks were conducted using both EEG and fMRI in separate sessions with matched task protocols and counterbalanced condition orders.

### Participants

A total of 58 healthy volunteers participated in this study. The cohort included 14 women and 44 men, with a mean age of 23.07 ± 1.99 years. Handedness was assessed using the Japanese version of the FLANDERS handedness questionnaire (Okubo et al., 2014), which yields scores ranging from -10 to +10, with larger positive values indicating stronger right-handedness. The mean handedness score across participants was 9.78 ± 0.77. Each participant completed two experimental sessions on separate days, one EEG session and one fMRI session, with the order counterbalanced across participants. The average interval between EEG and fMRI sessions was 17.72 ± 10.94 days. All participants were tested using an identical experimental protocol across sessions.

A written informed consent was obtained from all participants, who received monetary compensation. All recruitment procedures and experimental protocols were approved by the Institutional Review Board of Advanced Telecommunications Research Institute International (ATR, Kyoto, Japan) (approval number: 21-143). The study was conducted in accordance with the principles of the Declaration of Helsinki.

### MRI acquisition

Functional and structural MRI data were systematically acquired from all participants using a 3T Siemens MAGNETOM Prisma scanner at the ATR Brain Activation Imaging Center, equipped with a 32-channel head coil.

Functional images were obtained using a gradient-echo multiband echo-planar imaging sequence (multiband acceleration factor: 6) with the following parameters: repetition time (TR), 0.8 s; echo time (TE), 34 ms; flip angle, 52°; field of view, 206 × 206 mm; voxel size, 2.4 × 2.4 × 2.4 mm^3^; matrix dimensions, 86 × 86; and slice gap, 0 mm. Each run comprised 1160 volumes for the HCP-mini and 1106 volumes for the N-back. Two runs were performed for each of the HCP-mini and N-back tasks, resulting in a total of four runs per participant. For both tasks, images were acquired in an anterior-to-posterior (AP) phase-encoding direction in run 1 and posterior-to-anterior (PA) direction in run 2. For susceptibility distortion correction of the functional images, a spin-echo field map was acquired with the following parameters: TR, 6.1 s; TE, 60 ms; flip angle, 90°; field of view, 206 × 206 mm; matrix size, 86 × 86; voxel size, 2.4 × 2.4 × 2.4 mm^3^; and slice gap, 0 mm. Acquisition was repeated twice, and the second dataset was used for subsequent preprocessing.

Structural images were acquired as follows: T1-weighted images were obtained using a magnetisation-prepared rapid gradient-echo sequence (TR, 2.5 s; TE, 2.18 ms; inversion time (TI), 1000 ms; flip angle, 8°; voxel size, 0.8 × 0.8 × 0.8 mm^3^; field of view, 256 × 256 mm; matrix dimensions, 300 × 320; slice thickness, 0.8 mm; and slice gap, 0 mm), while T2-weighted images were acquired using a three-dimensional sequence (TR, 3.2 s; TE, 564 ms; flip angle, 120°; voxel size, 0.8 × 0.8 × 0.8 mm^3^; field of view, 256 × 256 mm; matrix dimensions, 300 × 320; slice thickness, 0.8 mm; and slice gap, 0 mm).

All data were converted from raw DICOM images to NIfTI files, the standard neuroimaging format (Neuroimaging Informatics Technology Initiative), using the software dcm2niix (Li et al., 2016).

### EEG acquisition

EEG data were recorded at a sampling rate of 1000 Hz using an EEG amplifier (BrainAmp MR; Brain Products GmbH, Gilching, Germany) in an electrically shielded EEG booth (PHT-AN-100; Shield Room Co., Ltd., Japan). A total of 32 electrodes were placed on the scalp, according to the international 10-10 system, at the following sites: Fp1, Fp2, Fz, F3, F4, F7, F8, F9, F10, FC1, FC2, FC5, FC6, Cz, C3, C4, T7, T8, CP1, CP2, CP5, CP6, Pz, P3, P4, P7, P8, P9, P10, Oz, O1, and O2. The ground and reference electrodes were positioned at Fpz and FCz, respectively, and the EEG signals were re-referenced offline to the average potential. Electrode impedances were maintained below 25 kΩ. Experimental event triggers were delivered to the EEG system via a dedicated trigger interface (TriggerBox Plus; Brain Products GmbH, Gilching, Germany), ensuring precise synchronisation of stimuli and task events with EEG recordings.

### Experimental setup

In the EEG experiment, the participants were comfortably seated in an electrically shielded booth facing a 24-inch colour LCD monitor (EV2456; EIZO Co., Ltd., Japan) at a distance of approximately 57 cm. The head position was stabilised using a chin rest to minimise movement-related noise, with the gaze naturally oriented toward the centre of the screen. Responses were recorded using two handheld buttons (CrazySmall OneFT; cooyou.org) in the participants’ left and right hands. Visual stimuli were presented at a resolution of 1024 × 768 pixels for the HCP-mini task and 1920 × 1080 pixels for the N-back task. For the language task, auditory narratives were delivered through the monitor’s audio output.

In the fMRI experiment, the participants lay supine on the MRI bed and viewed the stimuli presented on a monitor inside the scanner bore via a mirror mounted on the head coil. The tasks were performed while the participants held an MRI-compatible response button box (Bimanual 4-Button Fibre Optic Response Pad; Current Designs, Inc., Philadelphia, PA, USA) in both hands. The viewing distance was approximately 1080–1095 mm. Visual stimuli were presented at a resolution of 1024 × 768 pixels for the HCP-mini task and 1920 × 1080 pixels for the N-back task, consistent with the EEG experiment. For the language task in the fMRI session, auditory narratives were presented using MRI-compatible headphones where head-coil fit allowed. Of the 58 participants, 32 received binaural presentation and 25 received monaural presentation to the right ear because bilateral headphone placement was constrained by head-coil fit. In the remaining participant, headphones could not be fitted.

### Procedure

#### HCP-mini task

This task is an integrated behavioural paradigm designed to assess multiple cognitive domains within a single run, based on the HCP task design principles (Barch et al., 2013). While maintaining conceptual and structural correspondence with the original HCP tasks, the paradigm was reconfigured to accommodate the time constraints associated with EEG and fMRI acquisition.

All tasks were adapted from the original HCP tasks programmed in E-Prime 2.0 and modified using E-Prime 3.0 (version 3.0.3.219; Psychology Software Tools). Each task had an approximate duration of 80 seconds, and eight cognitive tasks—emotion processing (EMO), social cognition (SOC), relational processing (REL), gambling (GAM), language (LAN), motor (MOT), picture-based working memory (WM-p), and letter-based working memory (additional task; WM-l)—were administered sequentially within the same run. A resting-state condition (REST) was included, resulting in nine task conditions per run (Figure 2). For tasks involving binary choice responses, the left–right position of the correct option on the screen was counterbalanced across participants to minimise systematic response bias.

**Figure 2.**
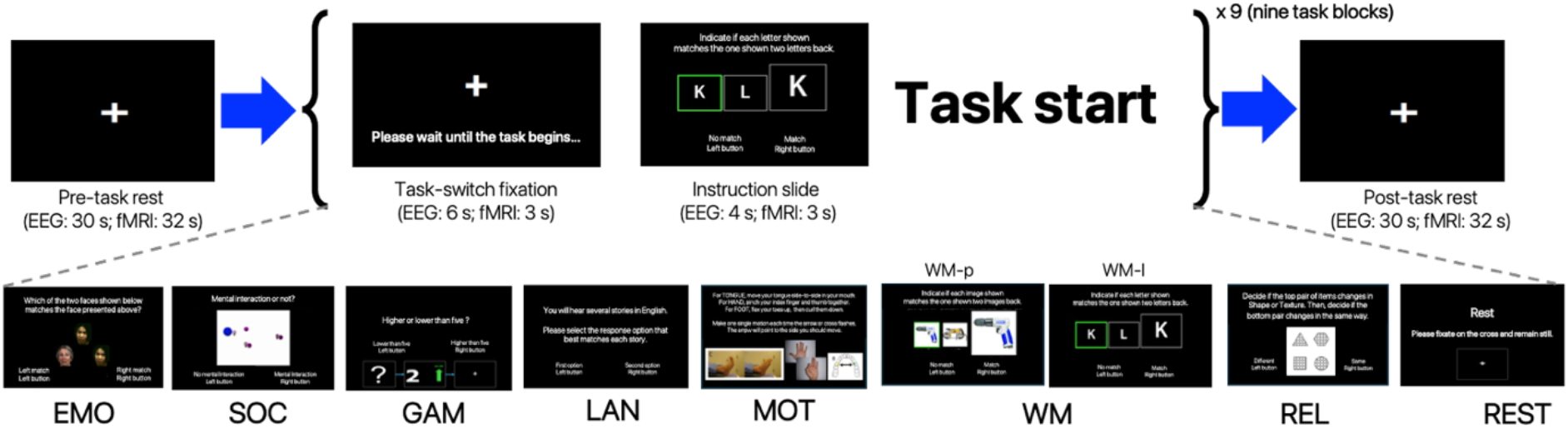
Schematic overview of the Human Connectome Project-aligned multitask paradigm (HCP)-mini task structure within a single run. Each run consists of nine task blocks, including eight cognitive tasks—emotion processing (EMO), social cognition (SOC), gambling (GAM), language (LAN), motor (MOT), picture-based working memory (WM-p), letter-based working memory (WM-l), and relational processing (REL)—and a resting-state condition (REST). At the beginning of each run, a 4-second countdown screen from the original HCP implementation is presented before the pre-task rest period. The task sequence is preceded by a pre-task rest, and before each task block, by a task-switch fixation and an instruction slide. The same structure is repeated across task blocks, and a post-task rest period is included at the end of the run.

Each task preserved the essential stimulus structure and trial characteristics of its corresponding HCP task while being temporally shortened to ensure that the full run could be completed within approximately 15 minutes. Tasks were presented continuously, enabling uninterrupted recording of behavioural responses and neural activity throughout the run. The order of task presentation was randomised across the participants to minimise the systematic effects of task order or fatigue. Stimulus presentation timings, trial onset and offset times, and participant responses were recorded and synchronised with EEG and fMRI data.

On both EEG and fMRI acquisition days, the participants completed a practice run for each task before the main experiment, following an identical procedure across modalities. Practice trials were conducted in a fixed order (EMO, SOC, GAM, LAN, MOT, WM-p, WM-l, REL). For tasks with objectively defined correct responses, practice continued until participants demonstrated sufficient understanding, operationalised as achieving an accuracy level of approximately 80%.

Given that the primary aim of the HCP-mini task was to enable comparison across specific cognitive functions within a limited acquisition time, the task design focused on experimental conditions, without including the explicit control conditions used in the original HCP tasks. For example, while the original HCP emotion processing task included both face and shape conditions, whereas the HCP-mini task included only the face condition. In the original HCP tasks, these control conditions were primarily used in fMRI-based general linear model analyses to compare the experimental and control conditions (e.g. face vs. shape) (Barch et al., 2013). In contrast, the HCP-mini task adopts a cross-task comparative design, in which the neural activity associated with a given task is evaluated relative to the average activity observed across all tasks within the same run. This design emphasises the identification of task-dominant neural activity profiles within the run as a whole, rather than relying on classical contrasts using explicit control conditions. Accordingly, rather than isolating task effects through dedicated control conditions, the present design characterised relative activation patterns across cognitive domains within a multitasking framework.

Thus, the HCP-mini task was designed to sample multiple cognitive domains within a single run while retaining a broad alignment with the HCP task taxonomy. Each task was adapted from the original HCP framework with temporal modifications and included an additional working memory task to enable integration within a compact multitask design (Figure 3).

**Figure 3.**
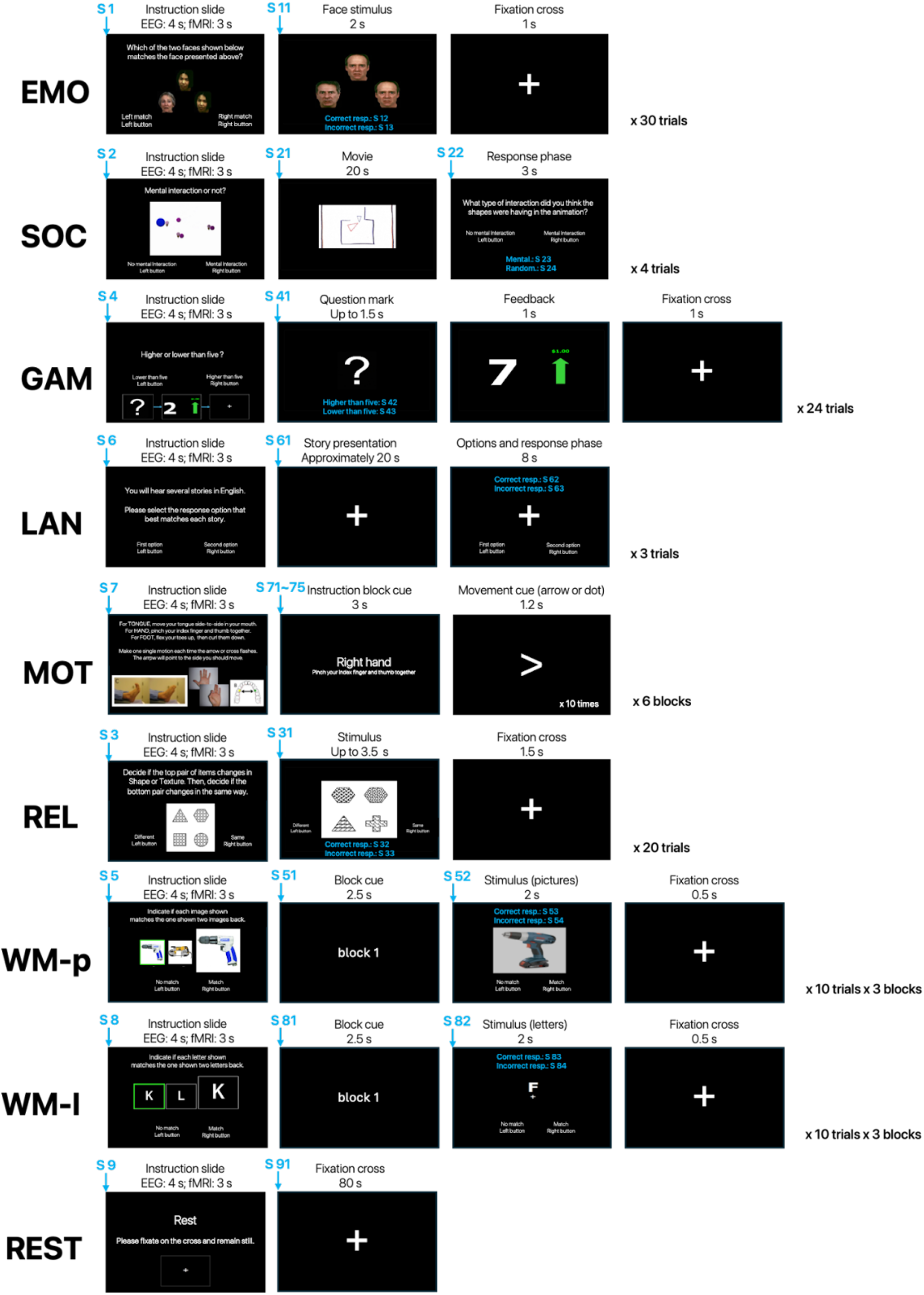
Trial structure of the Human Connectome Project-aligned multitask paradigm (HCP)-mini task. This figure illustrates the temporal sequence of events within a single trial for each task. Each row shows the order of the stimulus presentation and response phases for one representative trial of the corresponding task. The blue labels (S × x) indicated by arrows denote the EEG trigger codes sent at the onset of each event. The durations of individual events are shown in the corresponding panels. The number of trials or blocks per run is indicated on the right side of each task sequence. **Abbreviations** EMO, emotion processing; SOC, social cognition; GAM, gambling; LAN, language; MOT, motor; REL, relational processing; WM-p, picture-based working memory; WM-l, letter-based working memory; REST, resting-state condition.

#### Emotion processing

This task was based on the emotional processing paradigm used in the HCP, originally introduced by Hariri et al. (2002), and it comprises perceptual matching judgement using facial stimuli (anger or fear). In each trial, a single target face stimulus was presented at the top of the screen together with two face stimuli at the bottom, and the participants selected the stimulus that matched the target. Each face stimulus was presented for 2 seconds, followed by a 1-second inter-stimulus interval with a fixation cross. Each task consisted of 30 trials. A set of 12 face stimuli was used, which were presented repeatedly in a randomised order across trials.

#### Social cognition

This task was based on the social cognition (theory of mind) paradigm used in the HCP, developed in previous studies, in which animated stimuli depict social interactions between geometric shapes (Castelli et al., 2000; Wheatley et al., 2007). Animated video clips depicting social interactions between geometric shapes were presented to the participants. In each trial, a video clip lasting 20 seconds was presented, followed by a 3-second response interval, in which the participants indicated whether the animation depicted a social interaction. The task consisted of four randomly selected trials from a set of eight video clips. In contrast to the original social cognition task, which employed a three-option response format (social interaction, unsure, or no interaction), the present task used a two-choice response format to maintain consistency with the response structure adopted across other tasks in the HCP-mini paradigm. In addition, only video clips depicting social interactions were used.

#### Gambling

This task was based on the gambling (incentive processing) paradigm used in the HCP, which was developed by Delgado et al. (2000). In each trial, participants were presented with a question mark and required to predict whether the hidden value, ranging from 1 to 9, would be higher or lower than 5 using a two-choice button. The prediction phase lasted for up to 1.5 seconds; when a response was made earlier, a fixation cross was displayed for the remaining time. This was followed by a 1-second feedback phase, during which the actual value was revealed together with a visual outcome cue indicating a win, loss, or neutral outcome, depending on the participant’s prediction.

Each trial ended with a 1-second inter-stimulus interval with a fixation cross. Each task consisted of 24 trials. The trials were organised into three blocks of eight, with outcomes biased toward either reward or punishment. The three blocks were administered in a fixed order within each run, and no explicit block cues were presented. In contrast to the original gambling task, the fixation blocks were not included.

#### Language

This task was based on the language processing paradigm used in the HCP, which was adapted from the auditory story comprehension task developed by Binder et al. (2011). The participants listened to short auditory narrative stimuli and subsequently answered a comprehension question. In each trial, an auditory story lasting approximately 20 seconds was presented, followed by an 8-second response interval, during which participants selected the correct answer from two alternatives using buttons. The task consisted of three trials randomly selected from a pool of nine narrative stimuli, which were selected from a larger set.

#### Motor

This task was based on the motor-mapping paradigm used in the HCP, which was developed in previous studies (Buckner et al., 2011; Yeo et al., 2011). The participants performed visually cued movements involving different effectors, including the hands, feet, and tongue. Specifically, they tap their left or right fingers, squeeze their left or right toes, or move their tongues. Each movement block consisted of a brief cue (3 seconds) followed by a 12-second period of repetitive movement execution (10 repetitions). Six movement blocks were conducted, and the order of the movement types varied between runs.

#### Relational processing

This task was based on the relational processing paradigm used in the HCP, which was adapted from the relational reasoning task developed by Smith et al. (2007). Participants were presented with two pairs of objects; one pair displayed at the top of the screen and the other at the bottom. They were instructed to first determine which dimension differed between the objects in the top pair (shape or texture), then judge whether the objects in the bottom pair also differed along the same dimension (e.g. if the top pair differed in shape, whether the bottom pair also differed in shape). In each trial, the stimulus was presented for up to 3.5 seconds within a fixed trial duration of 5 seconds. If a response occurred within 3.5 seconds, a fixation cross was displayed for the remainder of the trial. The task consisted of 20 trials, with stimulus combinations randomly selected from a set of 48 possible stimuli. In contrast to the original relational processing task, the control matching condition was not included in the present task.

#### Picture-based working memory

This task was based on the working memory paradigm used in the HCP, which was adopted because it could serve as a functional localiser of working memory (Drobyshevsky et al., 2006). In the task, the stimuli consisted of four categories: faces, places, tools, and bodies. Each trial consisted of a 2-second stimulus presentation followed by a 0.5-second inter-stimulus interval, during which a fixation cross was displayed. A cue (2.5 seconds) was presented at the beginning of each block. The task consisted of three blocks with a total of 30 trials. The trial order within each block was fixed, whereas the stimulus sets varied between runs.

#### Letter-based working memory

This task employed an N-back working memory design analogous to the picture-based working memory task using alphabetical letter stimuli, based on the working memory paradigm used in the HCP. Each trial consisted of a 2-second stimulus presentation followed by a 0.5-second inter-stimulus interval, during which a fixation cross was displayed. A cue (2.5 seconds) was presented at the beginning of each block. The task consisted of three blocks with a total of 30 trials. The trial order within each block was fixed, whereas the stimulus sets varied between runs.

#### Resting-state

During the resting-state condition, participants did not perform any explicit tasks and remained at rest for 80 seconds. They were instructed to relax (while looking at a fixation cross presented at the centre of the screen), remain awake, and not think about anything.

### N-back task

The participants performed a visual N-back working memory task designed to systematically vary the cognitive load within a single-task framework (Figure 4). The task consisted of eight conditions (0-back to 7-back) and one 80-second resting block. Unlike the task battery derived from the HCP, which is designed to enable comparisons across distinct cognitive tasks, this N-back task enables within-task comparisons across conditions by parametrically manipulating the working memory load. In this study, the N-back task was acquired as a separate task battery from the HCP-mini battery, enabling comparisons across N-back levels, as well as broader comparisons with the HCP-derived tasks.

**Figure 4:**
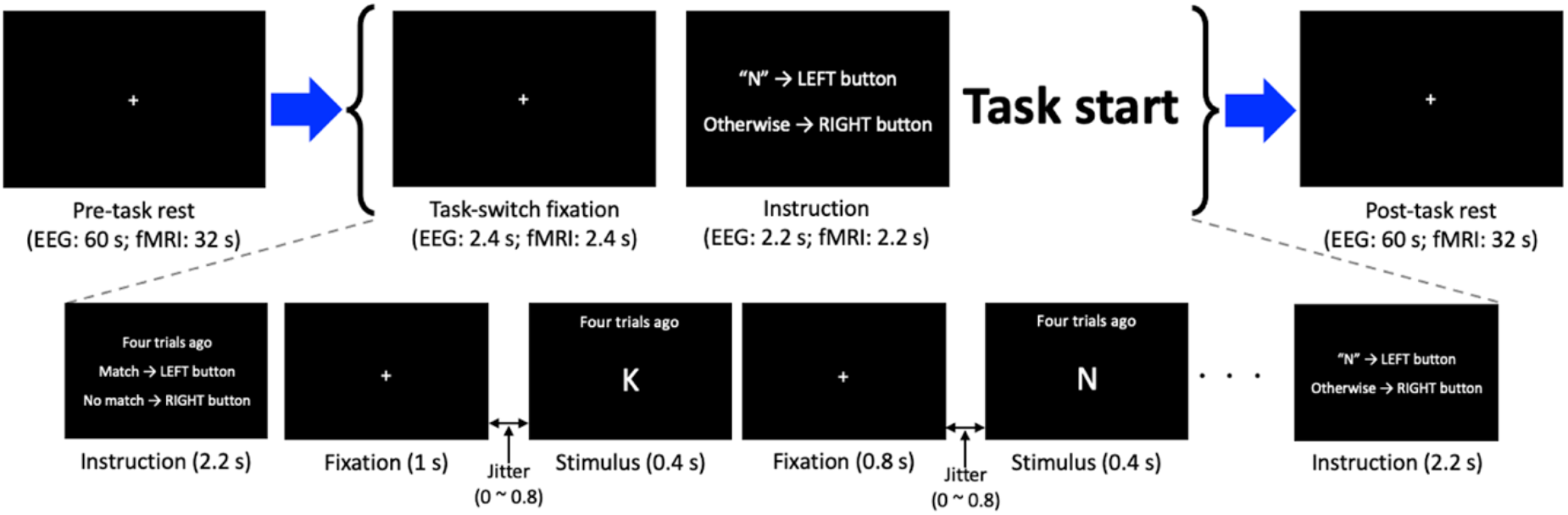
Experimental design of the extended N-back task. The N-back paradigm consists of a pre-task rest, task-switch fixation, instruction, and task periods, followed by a post-task rest. During the task period, participants perform a continuous N-back task in which they respond to each stimulus based on whether it matches the item presented in the *N* trials earlier. Each trial consists of a stimulus presentation (0.4 seconds) interleaved with fixation periods and temporal jitter (0.8 seconds). Instructions are presented at the beginning of each block to indicate the task rule (e.g. match vs. non-match). The figure illustrates an example sequence (shown here for the 4-back condition and 0-back conditions). Timing parameters are shown for both EEG and fMRI data. Abbreviations: EEG, electroencephalography; fMRI, functional magnetic resonance imaging

Each recording session consisted of an initial resting period, followed by nine conditions (eight N-back blocks corresponding to the 0-7-back conditions and one rest block) in randomised order, and ended with a final resting period. In the EEG session, participants first completed a 60-second resting period, followed by nine conditions, concluding with a final 60-second resting period. Short fixation periods (“task-switch fixation”) were interleaved between N-back blocks. In the fMRI session, the same structure was used, except that the initial and final resting periods were 32 seconds each.

Single uppercase letters were used as stimuli. For each session, a subset of nine letters was selected randomly from the English alphabet and used throughout the session. The selected letter sets differed across sessions and among participants.

Each task block began with an instruction period (2.2 seconds), followed by a fixation period (1.0 second), after which a sequence of trials for the block commenced. During the fixation periods, participants were instructed to focus on the central fixation marker without responding.

On each trial, stimulus onset was preceded by a variable pre-stimulus interval (jitter), followed by presentation of a single uppercase letter for 0.4 seconds, and a fixed post-stimulus fixation period of 0.8 seconds (Figure 4). The pre-stimulus interval had a mean duration of 0.4 seconds, resulting in a mean trial duration of 1.6 seconds. A fixation marker consisting of a double circle with a filled inner circle was continuously displayed throughout the task, including during stimulus presentation, serving as a visual reference across trials.

Each N-back condition consisted of 50 + *N* trials. Target trials accounted for 30% of trials (15 trials per block) and lure trials accounted for 20% (10 trials per block). Lure trials were implemented only for tasks with *N* ≥ 2. When enabled, lure trials were defined as trials in which the current stimulus matched the stimulus presented at the *N*−*1* or *N+1* positions, while excluding exact *N*-back matches. For the 2-back task, lure trials were included only in a subset of blocks, as indicated by the event annotations. Target stimuli were not presented during the first *N* trials of each block.

Participants responded to every trial using a two-alternative forced-choice (2AFC) button press, indicating whether the current stimulus matched the N-back criterion. In the 0-back condition, participants indicated whether the currently presented stimulus matched a predefined target stimulus. The assignment of response buttons (left/right) to the target and non-target responses was counterbalanced across participants.

The EEG and fMRI data were obtained on separate days. The task instructions, stimulus parameters, block structure, and response mappings were identical across modalities to minimise session-dependent variability. Before data acquisition, the participants completed a practice session. Practice continued until performance in the 0-back and 1-back conditions exceeded 80%, defined as *“hit rate* − *false alarm rate”*. The first practice round included the 0-, 1-, 2-, 4-, and 6-back conditions, whereas subsequent practice rounds included only the 0-, 1-, and 2-back conditions. Each practice block consisted of 20 trials (6 target and 4 lure trials).

## Data analysis

### MRI preprocessing

The fMRI data were preprocessed using a combination of FSL and SPM12. First, susceptibility-induced distortion correction was performed using FSL’s TOPUP algorithm with pairs of images obtained in reversed phase-encoding directions. The estimated field maps were then applied to correct the geometric distortions in the echo-planar imaging (EPI) data.

Subsequent preprocessing was conducted using SPM12 (Wellcome Trust Center for Neuroimaging, London, UK). Functional images were realigned to the mean functional image to correct for head motion, and six motion parameters were estimated. The mean functional image was coregistered with individual structural (T1-weighted) images. The structural images were segmented and spatially normalised to the Montreal Neurological Institute (MNI) space, and the resulting deformation fields were applied to the functional images. Finally, the normalised functional images were spatially smoothed using a Gaussian kernel (full width at half maximum = 6 mm).

### Task activation analysis

A first-level statistical analysis was performed using SPM12 within the framework of a general linear model. For each participant, the task conditions were modelled as a boxcar function, with durations corresponding to the trial length, and convolved with a canonical haemodynamic response function. For the HCP-mini paradigm, condition-specific effects were estimated relative to an implicit baseline, resulting in contrasting images representing task-related activation for each condition. Six head motion parameters estimated during realignment were included as nuisance regressors. A high-pass filter (cutoff = 256 s) and AR(1) model were applied to account for low-frequency drifts and temporal autocorrelation, respectively.

Group-level analyses were conducted to characterise task-related activation patterns across cognitive domains. For the HCP-mini paradigm, the contrast images obtained at the first level were entered into a second-level analysis. To enable cross-domain comparisons, we computed the mean activation across all task conditions at the group level, and evaluated each task relative to the cross-task mean. Specifically, for each task, activation maps were expressed as deviations from the mean activation across seven HCP-compatible task conditions (EMO, SOC, GAM, LAN, MOT, REL, and WM-p) within the same dataset. This approach allowed us to characterise domain-dominant and domain-general activation patterns within a unified multitasking framework. This approach avoids reliance on explicit control conditions and instead defines task selectivity relative to the distribution of activities across tasks (King et al., 2019).

To enable cross-dataset comparison, statistical maps (Z-maps) were obtained from a NeuroVault collection (collection 457; https://identifiers.org/neurovault.collection:457), a public repository for sharing unthresholded statistical maps (Gorgolewski et al., 2015). The deposited maps were based on previously published studies (Van Essen et al., 2013). These maps were processed using an analogous normalisation procedure. Specifically, for each dataset (HCP-mini and NeuroVault collection 457), the mean Z-map across all task conditions was computed and subtracted from the task map. The resulting maps were then standardised (z-score) to facilitate comparisons across datasets.

### Behavioural summary measures

To characterise behavioural performance across the recording contexts, we first derived session-level summary measures for each participant for the EEG and fMRI sessions separately. Each modality comprised two runs, and session-level scores were calculated by averaging the two runs within each modality. For the HCP-mini task, mean accuracy and mean reaction time (RT) were computed for each task. Notably, for the language task, RT was calculated from the onset of the first option. For the N-back task, mean discrimination sensitivity (d′) and mean RT were computed for each load level.

For RT analyses, anticipatory responses and extreme outliers were excluded. Specifically, for the HCP-mini task, responses faster than 200 ms were excluded from analysis. For the N-back task, trials with RTs shorter than 150 ms or longer than the participant-specific mean plus 2.5 standard deviations were excluded.

Intraclass correlation coefficients (ICCs) were computed for both between-modality (EEG vs. fMRI) and within-modality run-to-run comparisons. For between-modality analyses, one session-level behavioural score from the EEG session was compared with the corresponding score from the fMRI session for each participant. For within-modality analyses, run 1 and run 2 summary measures were compared separately within EEG and within fMRI. Because behavioural summary measures were derived from different numbers of trials across tasks, ICC estimates may also reflect differences in task structure. ICCs were estimated using a two-way mixed-effects, single-measurement, consistency model, as the two measurement contexts constituted a fixed pair. The primary aim was to assess the consistency of scores between EEG and fMRI and between run 1 and run 2, rather than their linear association (Bland & Altman, 1986). ICCs were calculated in R (version 4.5.2; R Core Team, 2025) using functions from the base psych package and reported with 95% confidence intervals (CIs), following standard recommendations for ICC-based reliability analyses (Koo & Li, 2016). Pearson’s correlation coefficients were also reported to aid interpretation.

For task performance in the HCP-mini, one-sample t-tests against a chance level of 0.5 were conducted for EMO and REL, where chance levels could be defined, with Holm correction applied for multiple comparisons. No such test was performed for SOC, because the variable of interest was the proportion of trials in which participants selected the mental rather than random interactions, which is not a conventional accuracy measure. LAN accuracy was not compared against chance because the number of trials per run was small (three trials per run). WM-p and WM-l were also not subjected to chance-level comparisons, as accuracy in these tasks is influenced by factors such as target prevalence, and the small number of target trials renders indices such as d′ unstable and unsuitable.

For the N-back task, discrimination sensitivity (d′) was compared against 0 at each load level in both modalities.

## Results

### Behavioural results

#### Behavioural performance and reaction times in HCP-mini and N-back tasks

Behavioural performance and RTs for the HCP-mini and N-back tasks in the EEG and fMRI sessions are shown in Figure 5.

**Figure 5.**
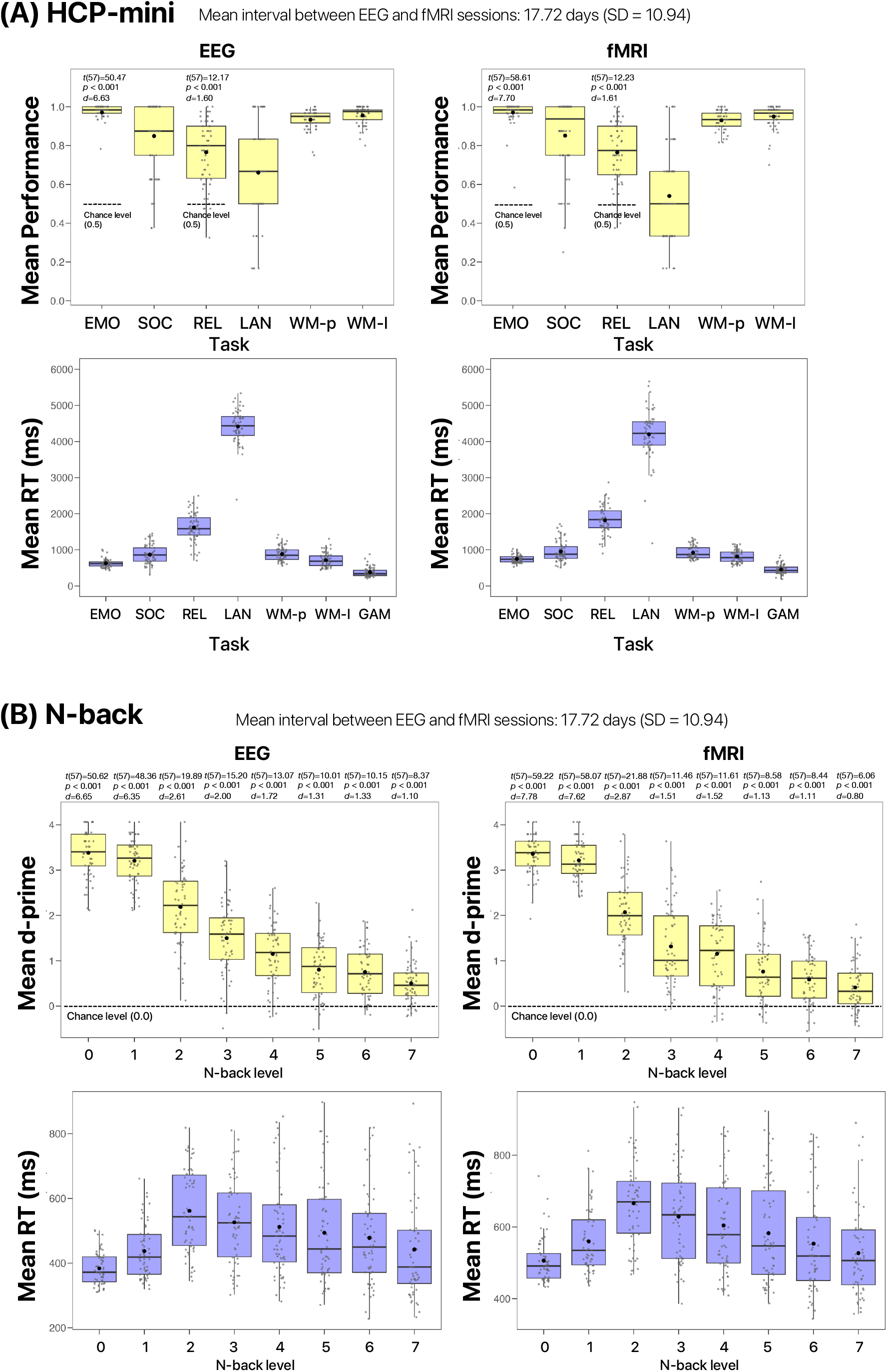
Behavioural performance and reaction time in the HCP-mini and N-back tasks. Human Connectome Project-aligned multitask paradigm (HCP)-mini task. Task-wise mean performance and mean reaction time (RT) are shown separately for the EEG and fMRI sessions. Emotion processing (EMO), relational processing (REL), language (LAN), working memory-picture (WM-p), and working memory-letter (WM-l) are shown as mean accuracy, whereas social cognition (SOC) is shown as the proportion of trials in which participants selected the mental rather than random interpretation. For EMO and REL, the horizontal line indicates the reference value of 0.5, and the corresponding statistical results are displayed (t value, Holm-adjusted p value, and Cohen’s d). (B) N-back task. Load-wise mean discrimination sensitivity (d′) and mean RT are shown separately for the EEG and fMRI sessions. For the d′ panels, the horizontal line indicates the reference value of 0, and the corresponding statistical results are shown in the figure (t value, Holm-adjusted p value, and Cohen’s d). Box plots indicate the median and interquartile range, whiskers indicate the range excluding outliers, and dots represent individual participants. Black dots indicate the mean.

In the HCP-mini, the mean accuracy is shown in Figure 5A (upper panel), and performance varied across tasks. Accuracy was highest for EMO and the two working memory variants (WM-p and WM-l), whereas broader inter-individual variability was observed for REL and lower performance for LAN. In SOC, performance was indexed not as accuracy but as the proportion of trials in which participants selected mental rather than random interpretation; this measure also showed relatively broad variability across participants. Similarly, RT varied systematically across tasks, with slower responses in REL and LAN tasks than in the other HCP-mini tasks.

Chance-level comparisons are shown in Figure 5 where applicable. For HCP-mini, mean accuracy for EMO and REL was significantly above 0.5 in both modalities. In contrast, SOC mental-response proportions, LAN accuracy, and WM-p and WM-l accuracy are shown descriptively only. For the N-back task, discrimination sensitivity (d′) was significantly above zero across load levels in both modalities.

The behavioural profile observed in the HCP-mini task was comparable to that reported in the original HCP framework. In the original HCP framework, qualitative comparisons were necessarily limited to tfMRI tasks for which behavioural accuracy distributions were presented (Barch et al., 2013, Figure S1), namely, EMO, LAN, REL, and WM task. Within this subset, EMO showed near-ceiling accuracy with most participants clustered at 0.97–1.00. LAN also showed high accuracy, with the story condition centered at approximately 0.80–1.00, alongside a lower outlying value near 0.50. In REL, accuracy was distributed more broadly, spanning 0.40–0.95. In WM, accuracy in the 2-back condition was centred at 0.80–0.85, with a broad distribution extending from 0.60 to 0.95. These qualitative similarities were observed in HCP-mini despite differences in task duration, task composition, and the exclusion of explicit control conditions (Figure 5A).

In the N-back task (Figure 5B), behavioural performance systematically declined with increasing working-memory load. Discrimination sensitivity (d′) decreased monotonically as load increased. RT increased from lower to intermediate load levels and gradually decreased at higher loads, with similar patterns observed in both EEG and fMRI sessions. Across load levels, RTs were consistently longer in the fMRI session than in the EEG session by approximately 100 ms, while preserving the same overall load-dependent profile.

#### Between- and Within-modality ICCs for behavioural measures

Figures 6 and 7 present scatter plots of accuracy for HCP-mini, d′ for N-back, and RT for both task batteries. The between-modality ICCs for the behavioural measures (accuracy, d′, and RT) are shown in Figure 6 for both the HCP-mini and N-back tasks, together with the corresponding 95% confidence intervals for the ICC estimates. Pearson’s correlation coefficients are also shown in the figure. Across measures, ICCs were numerically close to the corresponding Pearson’s correlation coefficients. In the HCP-mini task, between-modality ICCs varied across tasks for both accuracy and RT.

**Figure 6.**
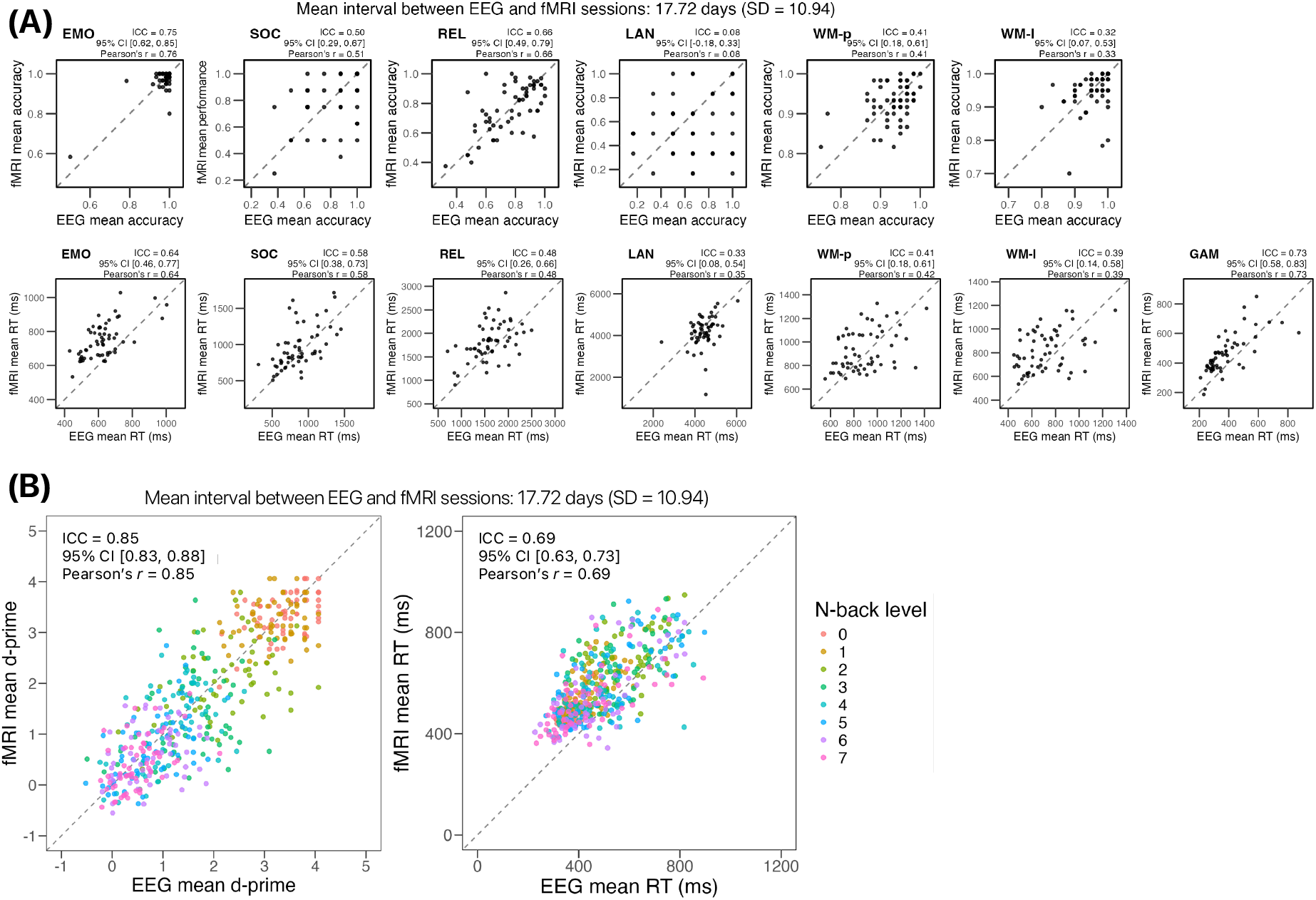
Between-modality ICCs for behavioural measures. (A)HCP-mini task performance, shown as mean accuracy (upper row) and mean reaction time (RT; lower row) for each task. (B)Pooled N-back performance across all load levels, shown for discrimination sensitivity (d′) (left) and RT (right), with points colour-coded by N-back level. ICCs and Pearson’s correlation coefficients are displayed in each panel. Each point represents one participant, and dashed lines indicate the line of identity. Axis ranges were adjusted separately across panels to preserve the visibility of the observed distributions. **Abbreviations** EEG, electroencephalography; fMRI, functional magnetic resonance imaging; HCP, Human Connectome Project; HCP-mini, Human Connectome Project-aligned multitask paradigm; EMO, emotion processing; SOC, social cognition; GAM, gambling; LAN, language; MOT, motor; REL, relational processing; WM-p, picture-based working memory; WM-l, letter-based working memory; REST, resting-state condition; ICC, intraclass correlation coefficient.

**Figure 7.**
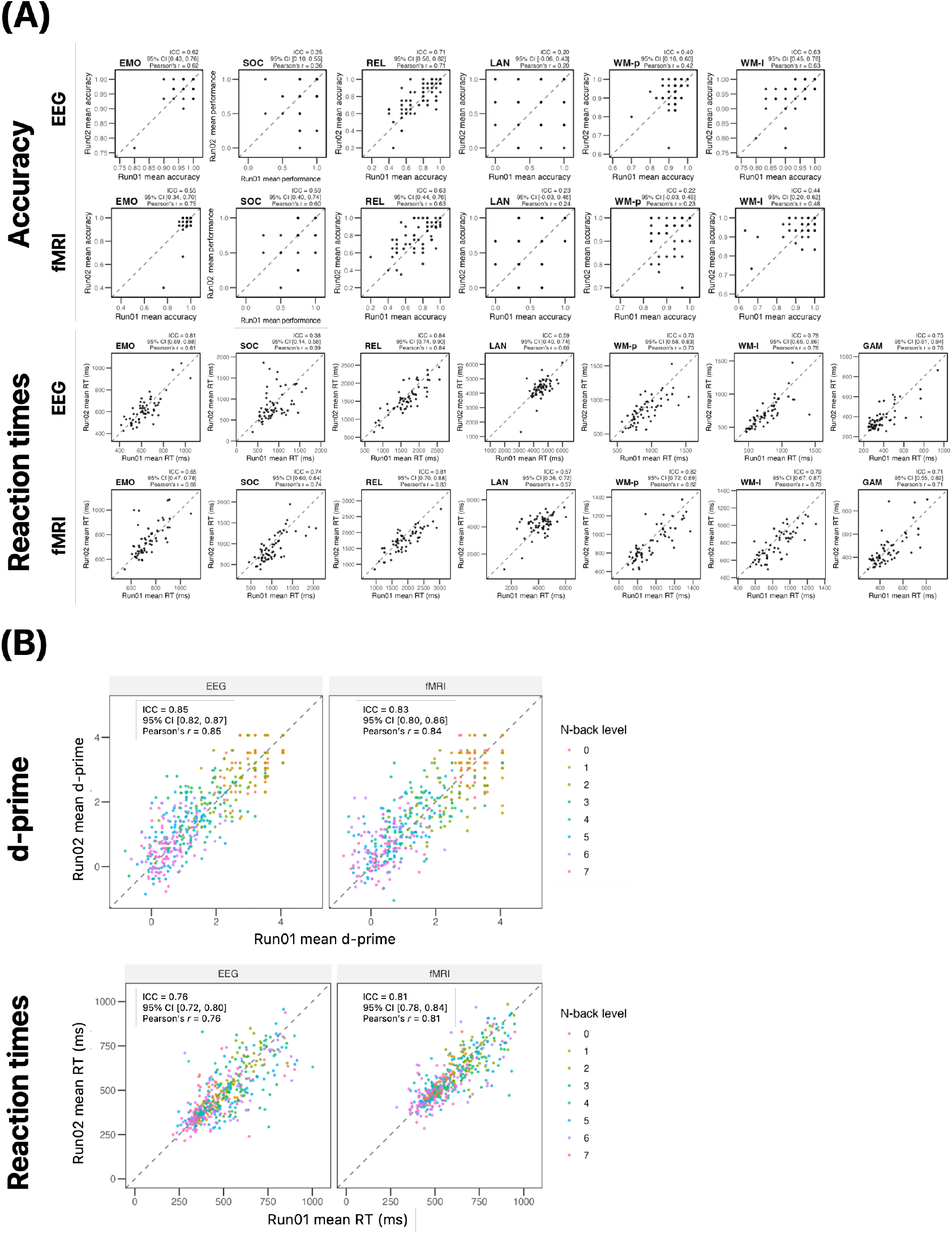
Within-modality run-to-run ICCs for behavioural measures. (A)HCP-mini task performance, shown as mean accuracy (upper row) and mean reaction time (RT; lower row) for each task. (B)Pooled N-back performance across all load levels, shown for discrimination sensitivity (d′) (upper row) and RT (lower row), with points colour-coded by N-level. ICCs and Pearson’s correlation coefficients are displayed in each panel. Each point represents one participant, and dashed lines indicate the line of identity. Axis ranges were adjusted separately across panels to preserve visibility of the observed distributions. **Abbreviations:** HCP-mini, Human Connectome Project-aligned multitask paradigm; ICC, intraclass correlation coefficient

The within-modality run-to-run ICCs for the behavioural measures are shown in the Figure 7 for both the HCP-mini and N-back tasks, along with the corresponding 95% confidence intervals for the ICC estimates. As in Figure 6, ICCs were numerically close to the corresponding Pearson’s correlation coefficients. In the HCP-mini task, within-modality ICCs also varied across tasks for both accuracy and RT.

Comparing Figures 6A and 7A, and Figures 6B and 7B, within-modality ICCs were generally numerically higher than the corresponding between-modality ICCs for both the HCP-mini and the pooled N-back measures.

#### fMRI activation map

To assess whether task-evoked activation patterns obtained from the HCP-mini dataset captured task-dependent functional organisation similar to that of the original HCP dataset, we compared spatial activation patterns. Visual inspection of the cortical activation maps revealed qualitatively similar patterns between the HCP-mini and HCP across all task conditions (Figure 8, left). Canonical large-scale networks associated with each task were consistently observed in both datasets despite differences in acquisition scale and sampling. To quantify this correspondence, we computed the spatial similarity between the task-evoked activation maps across the datasets using Pearson’s correlation. This analysis revealed a clear diagonal structure in the cross-dataset similarity matrix (Figure 8, right), with higher similarity for corresponding task pairs (e.g. WM–WM, motor–motor) than for non-corresponding pairs.

**Figure 8.**
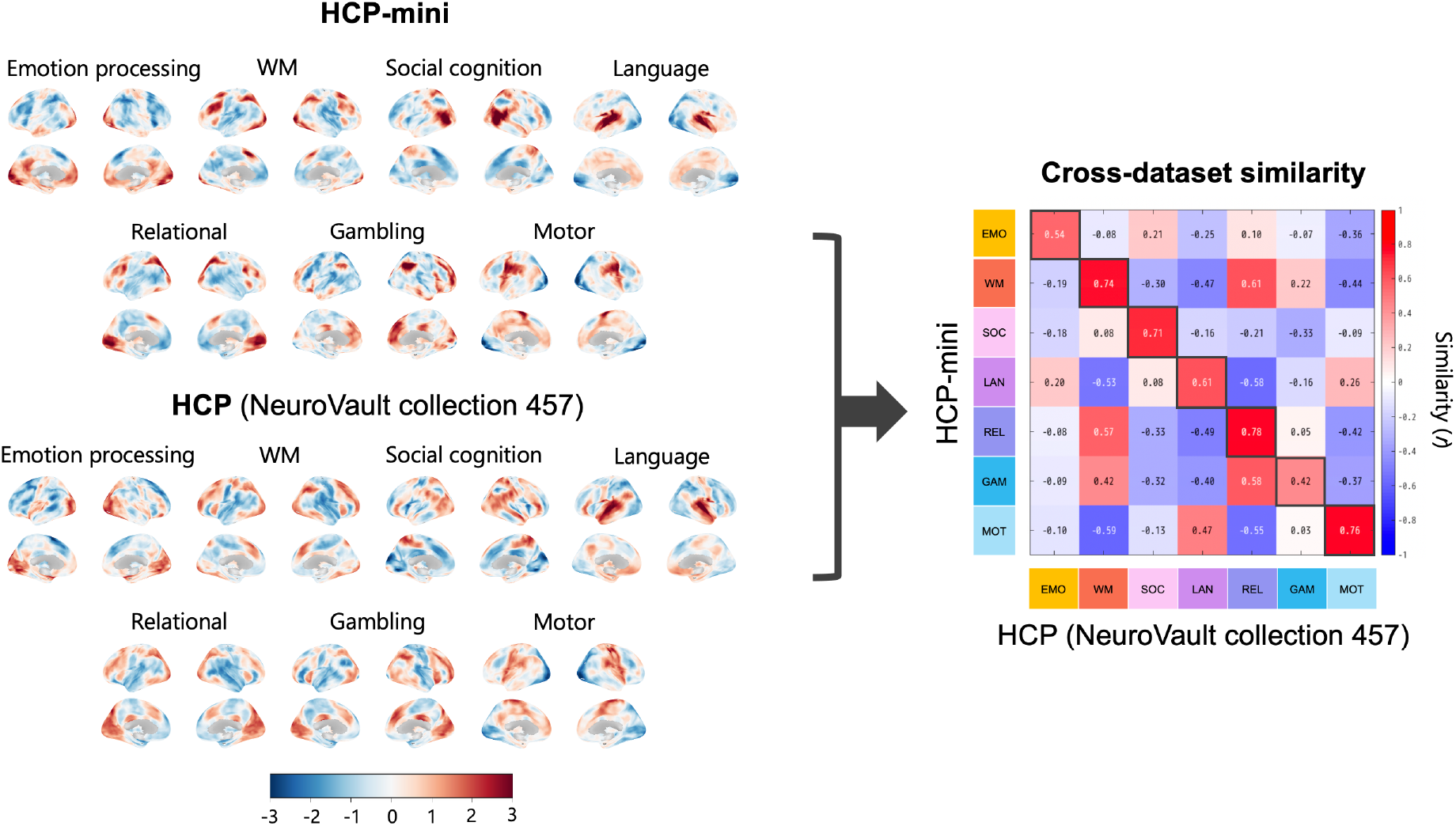
Cross-dataset similarity of task-evoked activation patterns. Spatial correspondence of task-evoked activation patterns between HCP-mini and HCP-derived statistical maps from NeuroVault (Collection 457). Left, cortical surface maps show task-evoked activation patterns for each task condition in the HCP-mini (top) and HCP (NeuroVault) datasets (bottom). Activation maps are displayed on inflated cortical surfaces (lateral and medial views) and represent normalised activation values (Z-scores). The task conditions include emotion processing (EMO), working memory (WM), social cognition (SOC), language (LAN), relational processing (REL), gambling (GAM), and motor (MOT). Right, cross-dataset similarity matrix quantifies the correspondence between task-evoked activation patterns across datasets. Each cell represents the spatial correlation (Pearson’s r) between the activation maps from the HCP-mini (rows) and HCP (columns). Diagonal elements indicate matched task pairs across datasets, whereas other elements reflect cross-task similarities. Higher similarity values along the diagonal demonstrate that task-specific activation patterns are preserved across datasets, indicating the robust cross-dataset reproducibility of large-scale functional organisation. **Abbreviations:** EEG, electroencephalography; fMRI, functional magnetic resonance imaging; HCP, Human Connectome Project; HCP-mini, Human Connectome Project-aligned multitask paradigm.

## Discussion

The present study examined whether a compact, within-run multitask design could preserve canonical behavioural structure and task-related functional organisation across cognitive domains, while providing a stable parametric manipulation of working-memory load across EEG and fMRI sessions. Overall, our findings support the functionality of this compact multimodal framework. The HCP-mini broadly preserved behavioural structure across cognitive domains and recovered broad cross-domain fMRI organisation consistent with the original HCP framework, whereas the N-back task provided a stable behavioural axis of graded cognitive demand.

### Behavioural findings in HCP-mini

The HCP-mini behavioural findings broadly supported the compact multitask design at the group level. The overall pattern across tasks was comparable to that reported for the original HCP framework (Barch et al., 2013; Van Essen et al., 2013), with near-ceiling performance for EMO, broader dispersion for REL, and lower performance for LAN. Among the HCP-mini tasks suitable for chance-level comparison, EMO and REL accuracy exceeded 0.5 in both modalities. However, SOC mental-response proportions, LAN accuracy, and WM-p and WM-l accuracy were interpreted descriptively rather than against chance, given the nature of these measures and the structure of the tasks. As aforementioned, with the exception of LAN, performance patterns were broadly comparable to those observed in the original HCP tasks, suggesting that the tasks achieved an acceptable degree of validity. In the N-back task, discrimination sensitivity (d′) exceeded zero at each load level in both modalities, suggesting that the task functioned appropriately.

Regarding ICCs, within-modality run-to-run ICCs were generally higher than the corresponding between-modality ICCs, and the ICC estimates were numerically close to the corresponding Pearson’s correlation coefficients. Given that between-modality comparisons spanned separate recording days, whereas the within-modality comparisons did not, this pattern is compatible with the influence of day-to-day variation and state-dependent factors on the between-modality estimates.

Overall, participants performed the tasks as intended, supporting interpretation of corresponding neural measurements in relation to the task conditions. However, these behavioural findings alone do not establish the validity or reliability of neural measurements.

Moreover, task-specific differences in ICCs should be interpreted cautiously because behavioural summary measures were based on different numbers of trials across tasks. This concern was especially relevant for SOC and LAN, which included only four and three trials, respectively. In addition, SOC was indexed as the proportion of “mental” responses rather than accuracy, limiting direct comparability with other measures. The relatively weak LAN ICCs likely reflect not only the very limited trial count and the use of English auditory materials in Japanese participants, but also modality-specific differences in auditory delivery, including monaural right-ear presentation in some fMRI participants and the absence of headphone presentation in one participant, as well as the susceptibility of auditory language processing in MRI to background noise and protocol-dependent variation (Hall et al., 1999; Peelle, 2014). These auditory-delivery differences may also have introduced additional variability in LAN-related neural responses, particularly in fMRI. Moreover, fMRI language mapping results vary substantially across speech comprehension protocols (Binder et al., 2008). Accordingly, the LAN results should be interpreted cautiously rather than taken to indicate a fundamental limitation of the task itself.

### Behavioural findings in N-back task

Discrimination sensitivity (d′) decreased monotonically with increasing load. Similarly, the RTs increased from lower to intermediate load levels and then gradually decreased at higher loads. These results are consistent with previous reports of nonlinear response profiles under increasing working memory demand (Lamichhane et al., 2020). In addition, d′ remained above zero across the analysed load levels, supporting the interpretation that the N-back task provided a behaviourally meaningful index of graded cognitive demand.

Within-modality ICCs for the N-back task were also numerically higher than the corresponding between-modality ICCs for both d′ and RT. As in the HCP-mini task, this pattern likely reflects the fact that between-modality indices incorporate influences beyond modality itself, including across-day effects and state-dependent variance. Unlike the HCP-mini analyses, however, N-back ICCs were calculated after pooling across N-levels; accordingly, these values reflect not only between- and within-modality relationships but also variance associated with task load. Moreover, RTs were longer in the fMRI session than in the EEG session, at least in part owing to differences in the response devices used (see Methods), a pattern also observed for the HCP-mini task. Nevertheless, the overall load-dependent pattern was preserved, suggesting that, as with d′, the task yielded behaviourally interpretable results.

Together, these findings indicate that behavioural performance under parametrically increasing working-memory load was relatively preserved, supporting the use of the N-back paradigm as a stable behavioural axis of cognitive demand. This, in turn, provides a useful behavioural foundation for examining neural activity associated with cognitive demand.

### fMRI activation map

The HCP-mini paradigm reproduced domain-specific activation patterns comparable to those observed in the HCP-derived statistical maps from the NeuroVault collection 457 (Van Essen et al., 2013). Thus, a compact multitask design can capture the core aspects of functional organisation across cognitive domains. By defining task-related activities relative to the mean across tasks, this approach captures relative functional selectivity rather than absolute activation levels, enabling direct comparisons across tasks and datasets without relying on explicit control conditions. Importantly, this framework requires the inclusion of a diverse set of tasks that broadly sample functional systems across the brain. Despite its compact design, the HCP-mini paradigm potentially provides sufficient coverage of cognitive domains to estimate the whole-brain functional organisation.

Motor-related tasks show consistent engagement of sensorimotor regions, whereas working memory tasks recruit frontoparietal networks, and language tasks engage distributed frontal and temporal areas (Owen et al., 2005; Price, 2012). These convergent patterns suggest that the relative activation framework preserves canonical functional organisation across datasets, which aligns with prior studies showing that large-scale brain organisation is preserved across task states (Yeo et al., 2011; King et al., 2019).

The clear diagonal structure observed in the cross-dataset similarity matrix quantitatively supports this connection, with higher similarity for the corresponding task conditions than for the mismatched pairs. This indicates that task-specific activation patterns are reproducible across datasets even when the acquisition scale and protocols differ. Some domains, such as GAM, showed weaker correspondence across datasets. This may reflect differences in task structure or variability in the engagement of reward-related processes; however, further investigation is required. Overall, these findings support the generalisability of this framework, indicating that relative task representations are preserved across datasets, despite differences in acquisition protocols and experimental designs.

### Limitations

This study had several limitations. First, the HCP-mini was developed as a compact adaptation of the original HCP tasks rather than as a direct replica. Task duration was shortened, some explicit control conditions were omitted, and certain tasks—particularly SOC and LAN—were implemented in a simplified form. Nevertheless, the behavioural and fMRI findings indicate that the compact design retained meaningful task-related structures across domains. Second, because EEG and fMRI were acquired on separate days, between-modality differences may reflect not only the recording context, but also state-dependent variation across sessions (Meltzer et al., 2007). Moreover, the number of trials contributing to each behavioural summary measure differed across tasks; therefore, the precision of these measures may vary across conditions. This should also be considered when interpreting task-specific differences. However, behavioural performance generally exceeded the relevant task-specific reference values, indicating that the task battery was appropriately established at the behavioural level for linkage with neural measurements in separate-session multimodal studies. Third, interpretation of the LAN findings warrants caution because the use of English materials by Japanese participants and modality-specific auditory delivery conditions may have introduced additional variability related to language comprehension. Thus, the weak between-modality ICC in this task should not be taken as evidence of a fundamental limitation of the task itself.

## Conclusions

This study demonstrated that a compact HCP-aligned multitask design can retain key comparative features of the original framework at both the behavioural and neural levels. In the HCP-mini task, group-level behavioural profiles were broadly preserved, and behavioural performance for the analysed measures generally exceeded the relevant reference values. In the N-back task, the expected load-dependent behavioural pattern was also observed across modalities, providing a stable behavioural axis of graded cognitive demand. Most importantly, the fMRI findings showed that integrating multiple cognitive tasks within a single run did not preclude recovery of canonical domain-related activation patterns that aligned with those of the original HCP dataset. Together, these findings support the usefulness of compact within-run multitask designs for multimodal studies of cognition across domains and graded cognitive demand.

## Acknowledgements

We thank Yuki Inoue for supporting the preparation and conduct of the experiment and Yoko Matsumoto for participant recruitment and scheduling. Furthermore, we would like to thank the team at the ATR Brain Activity Imaging Center (BAIC), in particular Akikazu Nishikido and Akihide Yamamoto, for their technical support. KH, SC, TA, MW, and AK were supported by the Innovative Science and Technology Initiative for Security (Grant Number JPJ004596), ATLA, Japan.

## Author contributions

KH designed and developed the experimental tasks, collected, analysed, and interpreted the data, and drafted the manuscript. SC contributed to task development, data collection, data analysis, data interpretation, and writing and revising the manuscript. AK and MW contributed to data collection. TA advised on the experimental implementation, data analysis, and interpretation. KH, SC, and TA contributed to the conception and design of the study under the supervision of HI. All authors read and approved the final manuscript.

## Competing interests

The authors declare no competing interests.

## Additional information

## Notes

### Competing Interest Statement

The authors have declared no competing interest.

